# The challenge of delimiting cryptic species, and a supervised machine learning solution

**DOI:** 10.1101/2021.08.05.455277

**Authors:** Shahan Derkarabetian, James Starrett, Marshal Hedin

**Affiliations:** Department of Organismic and Evolutionary Biology, Museum of Comparative Zoology, Harvard University, 26 Oxford St., Cambridge MA, 02138, USA; Department of Entomology and Nematology, University of California, Davis, Briggs Hall, Davis CA, 95616-5270, USA; Department of Biology, San Diego State University, 5500 Campanile Drive, San Diego CA, 92182-4614, USA

**Keywords:** integrative taxonomy, multispecies coalescent, RADSeq, short-range endemism, southern Appalachians, supervised machine learning, ultraconserved elements

## Abstract

The diversity of biological and ecological characteristics of organisms, and the underlying genetic patterns and processes of speciation, makes the development of universally applicable genetic species delimitation methods challenging. Many approaches, like those incorporating the multispecies coalescent, sometimes delimit populations and overestimate species numbers. This issue is exacerbated in taxa with inherently high population structure due to low dispersal ability, and in cryptic species resulting from nonecological speciation. These taxa present a conundrum when delimiting species: analyses rely heavily, if not entirely, on genetic data which over split species, while other lines of evidence lump. We showcase this conundrum in the harvester *Theromaster brunneus*, a low dispersal taxon with a wide geographic distribution and high potential for cryptic species. Integrating morphology, mitochondrial, and sub-genomic (double-digest RADSeq and ultraconserved elements) data, we find high discordance across analyses and data types in the number of inferred species, with further evidence that multispecies coalescent approaches over split. We demonstrate the power of a supervised machine learning approach in effectively delimiting cryptic species by creating a “custom” training dataset derived from a well-studied lineage with similar biological characteristics as *Theromaster*. This novel approach uses known taxa with particular biological characteristics to inform unknown taxa with similar characteristics, and uses modern computational tools ideally suited for species delimitation while also considering the biology and natural history of organisms to make more biologically informed species delimitation decisions. In principle, this approach is universally applicable for species delimitation of any taxon with genetic data, particularly for cryptic species.

## Introduction

Organismal diversity is underpinned by diversity in life history and ecological characteristics among taxa, which in turn produce different underlying genetic patterns at the population and species levels (Massatti and Knowles 2014, 2016; Fang et al. 2019; da Silva Ribeiro et al. 2020; Fenker et al. 2021). Biological characteristics can determine the process and type of speciation. For example, nonecological speciation (speciation without divergent natural selection) produces ecologically similar/identical species that are allo- or parapatric replacements of each other (Gittenberger 1991; Rundell and Price 2009) and is more likely in low dispersal taxa that also show niche conservatism (Czekanski-Moir and Rundell 2019). These biological and ecological characteristics often lead to cryptic speciation across many plant and animal taxa, where they prevent the application of many commonly used species criteria, and species delimitation relies largely, if not entirely, on genetic data.

The underlying diversity in speciation processes challenges the idea that any single genetic species delimitation method will be universally applicable. For example, the multispecies coalescent (MSC) is a popular model in genetic species delimitation, and many have argued for formalized delimitation under an MSC framework (Fujita et al. 2012). However, multiple empirical studies have shown that MSC models can over-split species level diversity in low dispersal taxa because such systems violate the underlying assumption of panmixia (Niemiller et al. 2012; Satler et al. 2013; Barley et al. 2013; Hedin et al. 2015; Hedin and McCormack 2017; Yang et al. 2019; Derkarabetian et al. 2019), a sentiment echoed in recent theoretical literature. Sukumaran and Knowles (2017) used simulations to show that BPP (Bayesian Phylogenetic and Phylogeography; Yang and Rannala 2010) equates population genetic structure with species level divergence. Barley et al. (2018) showed that model violation might bias inference of species boundaries in empirical “low dispersal” datasets, identifying subpopulations as species. These studies have not gone without rebuttals, focused on model implementation, simulation protocol, etc. (Leaché et al. 2019).

Regardless of ongoing debates, we argue simply that MSC implementations taken at face value, and with well-resolved genomic or sub-genomic data, have strong potential to inflate species numbers in low dispersal systems. A solution to over-reliance on genetics is integrative species delimitation (Dayrat 2005; Schlick-Steiner et al. 2010). However, in many poorly known or “cryptic biology” taxa (e.g., Jacobs et al. 2018), like minute animals that live under rocks and logs, integrating behavioral, ecological, and/or phenotypic data is challenging to impossible. Morphological conservatism resulting from niche conservatism (Wiens and Graham 2005) means that distinct species are often not morphologically diagnosable. In an integrative framework, some lines of evidence cannot be feasibly studied, while others are clearly conservative. This leads to a fundamental conundrum – how can we rigorously delimit species when genetic analyses are biased to inflate, and other evidence is inaccessible or lumps evolutionarily significant diversity?

Many taxa in the arachnid order Opiliones present challenges for species delimitation. Their microhabitat specificity and low dispersal ability leads to nonecological speciation and high population genetic structure, where related congeners rarely co-occur in sympatry ruling out direct tests for reproductive isolation (e.g., Thomas and Hedin 2008, Hedin and Thomas 2010; Derkarabetian et al. 2011; Derkarabetian and Hedin 2014, Starrett et al. 2016, DiDomenico and Hedin 2016, Derkarabetian et al. 2016; Peres et al. 2018; Derkarabetian et al. 2019). The biological characteristics associated with nonecological speciation in low dispersal Opiliones make several commonly used species criteria inapplicable or inappropriate in these systems. For example, morphological conservatism diminishes the utility of the morphological species criteria, and niche conservatism precludes ecological species criteria. In these cases, genetic data become the primary data type for species delimitation. However, these complexes represent classic cases of “too little gene flow”, where distinguishing population genetic structure from species level divergence is not easy, and genetic species delimitation analyses overestimate species diversity (e.g., Boyer et al. 2007; Fernández and Giribet 2014; Hedin et al. 2015; Derkarabetian et al. 2019).

Dispersal-limited microhabitat specialists are found in a diverse array of other taxa, including vertebrates and plants, with equal difficulty in resolving species-population boundaries (e.g., Niemiller et al. 2012; Yang et al. 2019). This under-appreciated issue remains one of the most difficult challenges for empirical species delimitation and its implications extend to a diverse array of taxa regardless of biological characteristics (e.g., Chambers and Hillis 2019). A possible solution to this issue is to use information from known systems to infer the unknown. In practice, this means using information derived from taxa with robust well-established species limits to infer species limits in a difficult cryptic species complex that shares similar biological and ecological characteristics and mode of speciation. Supervised machine learning is ideally suited for this problem, as known labeled data sets can be used to train a model that is then applied to an unknown unlabeled data set.

Here we use a combination of somatic and reproductive morphology, mitochondrial DNA, double-digest RADSeq (ddRAD; Peterson et al. 2012), and ultraconserved elements (UCEs; Faircloth et al. 2012; Starrett et al. 2017) to illustrate the species conundrum in *Theromaster brunneus* (Banks, 1902), a widely distributed species from the southern Appalachian Mountains with high potential for cryptic speciation (Fig. 1). Our goal is not to exhaustively test species limits using every data and analysis type, but instead to demonstrate the difficulty of delimiting species in such taxa using popular genetic species delimitation approaches, some of which are known to overestimate diversity. We highlight and emphasize a novel supervised machine learning approach, using training data from known taxa with similar biological characteristics as *T. brunneus*, to effectively delimit cryptic species using phylogenomic data.

**Figure 1.**
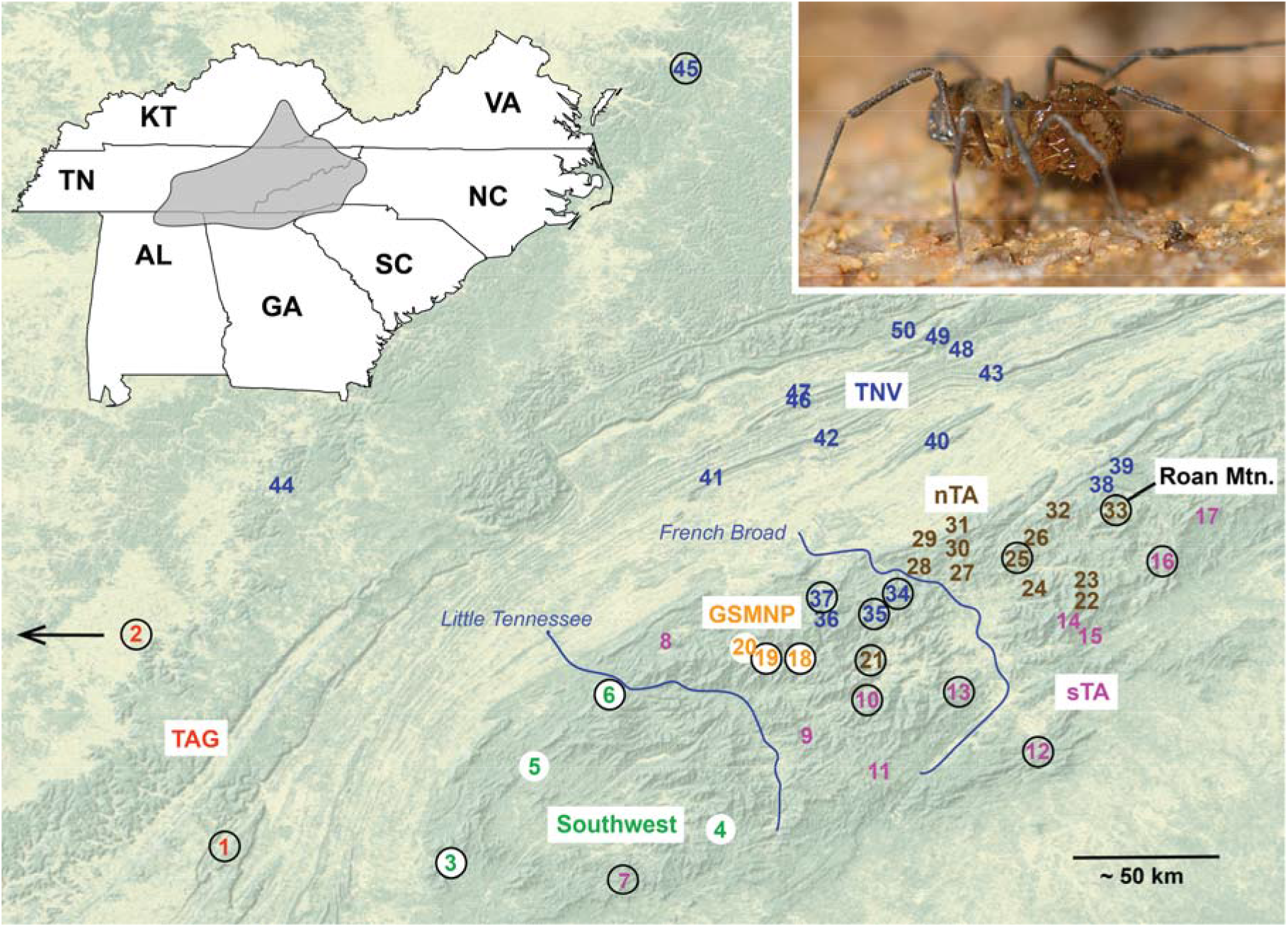
Distribution of *Theromaster brunneus* populations sampled in this study. Colors correspond to lineages identified in phylogenomic analyses. Numbers correspond to locations included in ddRAD analyses (Supplemental Table S1); sites with an enclosed circle were also sequenced for UCEs. Left inset: general distribution of *Theromaster*. Right inset: live photo of *T. brunneus*. Clade names: Great Smoky Mountain National Park (GSMNP), Tennessee Valley (TNV), northern Trans Asheville (nTA), southern Trans Asheville (sTA).

## New Approaches

We used a recently developed supervised machine learning approach for genetic species delimitation that was demonstrated with a “general” training data set based on a broad diversity of organisms. In this study, the full potential of a supervised machine learning approach is demonstrated by creating an empirical “custom” training data set, derived from a taxonomic group with robust well-defined species boundaries, and applying it to a cryptic species complex with similar biological characteristics. We operate under the premise that closely related taxa with similar biological and ecological characteristics and similar speciation processes should share similar underlying genetic patterns in regard to population- and species-level variation. In this way, the custom training data set becomes specific to this type of organism and data type, and our *a priori* knowledge of species in known systems is leveraged to infer species in unknown systems. When applied to a cryptic species complex the “custom” data set results in reasonable and biologically informed species hypotheses, as opposed to the “general” training data set, and other commonly used genetic species delimitation methods, which overestimate species numbers.

## Results

Voucher specimen data and accession numbers are provided in Supplemental Table S1. Raw ddRAD and UCE reads are available at the NCBI Short-Read Archive (BioProject [NNNN] and BioProject [NNNN], respectively), with matrices and tree files available from the Dryad Digital Repository: http://dx.doi.org/10.5061/dryad.[NNNN]. Representative morphological images (Fig. 2, Supplemental Figs S1-S5) confirm the highly conservative morphology across *Theromaster*, including male genitalia and male cheliceral modifications, a sexually dimorphic feature. However, there is clear differentiation in the habitus morphology of Roan Mountain (OP322, location #33), which has a more pointed eye mound, dorsal tergites clearly separated by grooves, more obvious dorsal spines, and more defined pigmentation patterns (Fig. 2). This specimen is clearly distinguished based on the morphological species criterion, suggesting at least two putative species. RAxML analysis of COI was mostly poorly supported but revealed highly divergent Roan Mountain and “Southwest” lineages; COI bPTP analyses supported 11 species (Supplemental Fig. S6).

**Figure 2.**
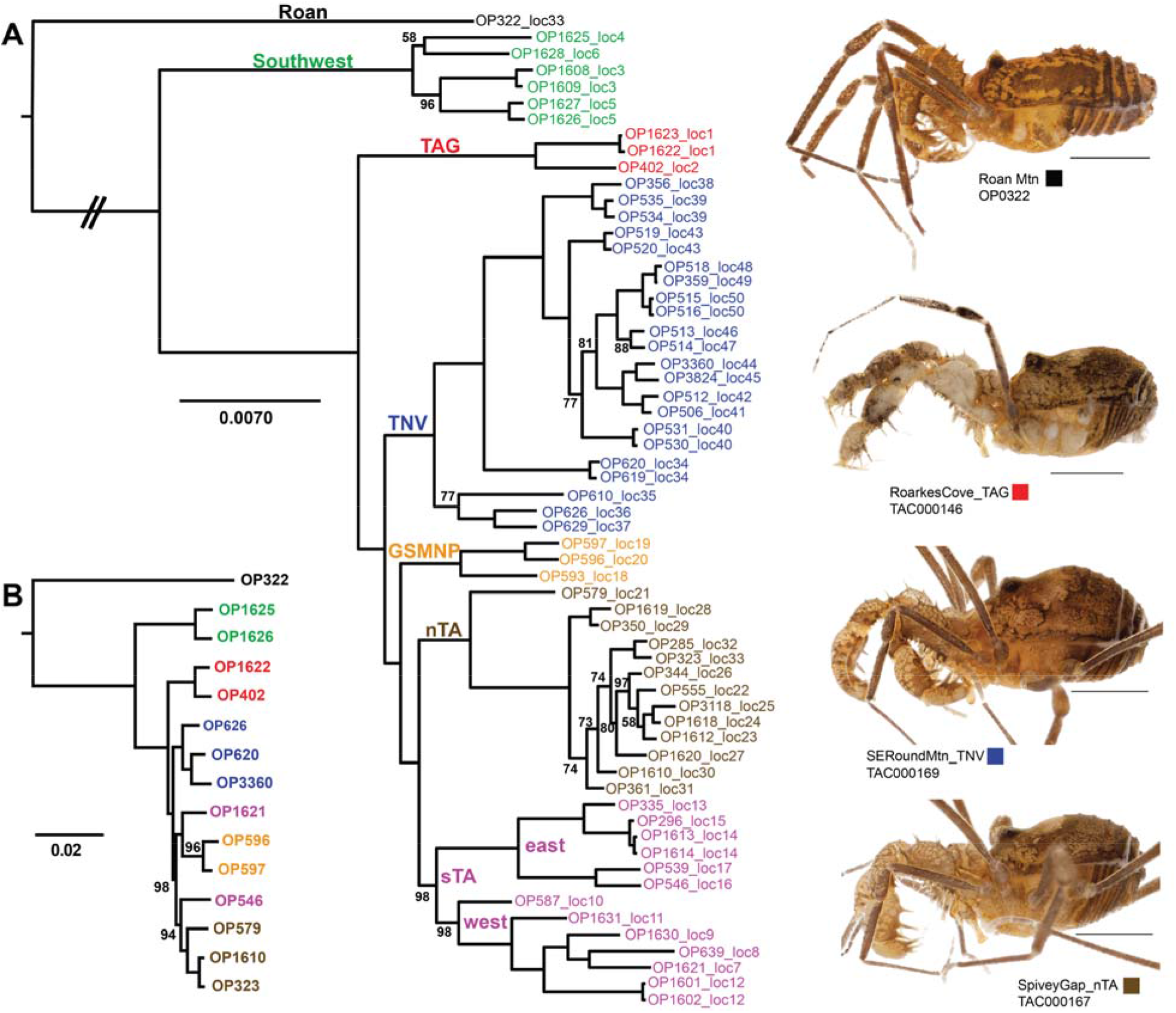
Phylogenomic results based on RAxML analysis of ddRAD and UCE data, with representative habitus images of select lineages. All nodes have 100% bootstrap support unless indicated. A) ddRAD phylogeny of the 61_48 dataset (see Supplemental Material I for matrix details). TNV, GSMNP, nTA, and sTA lineage samples were included in DAPC and STRUCTURE analyses. B) UCE RAxML phylogeny based on the 70% taxon occupancy matrix.

RAxML analyses of ddRAD data (∼75% taxon occupancy matrix with 1001 loci and 89,663 nucleotides) show at least seven main lineages (Fig. 2, Supplemental Fig. S7, Supplemental Tables S2-3). SVDQuartets analyses result in similar lineage composition and interrelationships, but with weaker support (Supplemental Fig. S8). Excluding a single instance of sympatry at Roan Mountain, all main lineages are allopatric (Fig. 1). Both partitioned RAxML and SVDQuartets analyses of the concatenated UCE matrix (70% taxon occupancy with 324 loci and 108,875 nucleotides) supports the ddRAD topology, with Roan Mountain, Southwest, and TAG as divergent and early-diverging lineages (Figure 2, Supplemental Fig. S9). DAPC clustering of the ddRAD SNPs reveal optimal K values (Supplemental Fig. S10) that correspond to the main lineages, with further subdivisions for the “southern Trans Asheville” (sTA) and “Tennessee Valley” (TNV) groups recovered in phylogenetic analyses. STRUCTURE analyses of ddRAD SNPs likewise recover the main lineages, but as K values increase, further subclusters are found that correspond to geographic and/or phylogenetic groupings (Supplemental Fig. S11). The best-fit K value is K = 3 using the ΔK method (Evanno et al. 2005), but K = 10 using the Pritchard et al. (2000) method. VAE analyses cluster samples as in other analyses, and UCEs show more overlap among groups (Supplemental Fig. S12).

BFD^*^ hypothesis testing results are presented in Figure 3 and Supplemental Tables S4-S5. Matrices consisted of 1828 and 415 SNPs for ddRAD and UCE datasets, respectively. The ddRAD dataset favored K = 3 corresponding to Roan Mountain, Southwest, and all other samples, while the UCE dataset favored all samples as species (K = 15), with a second less likely peak at K = 3 (Fig. 3). CLADES analyses based on the “general” training dataset favored all populations as species, while analyses based on the “custom” training dataset favored two cryptic species (Fig. 4).

**Figure 3.**
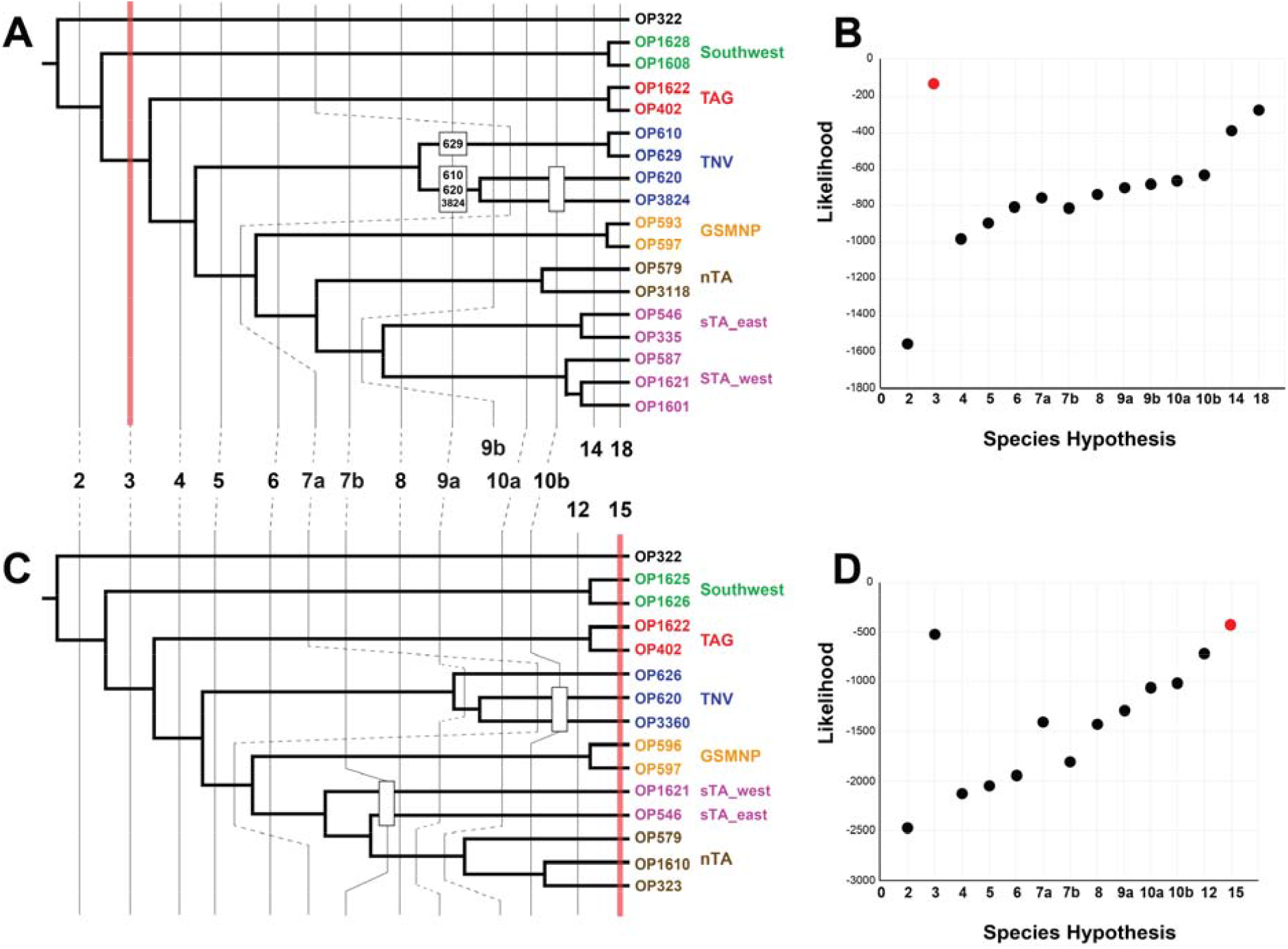
Results of SNAPP/BFD* species delimitation analyses based on SNP datasets. A and B) ddRAD phylogeny based on RAxML analysis of the 18_12 taxon occupancy matrix, with vertical lines indicating species hypotheses tested, and likelihood estimates for each hypothesis (averaged from two runs). C and D) UCE phylogeny based on RAxML analysis of the 70% taxon occupancy matrix, with vertical lines indicating species hypotheses tested, and likelihood estimates for each hypothesis (averaged from two runs). Note: branch lengths for both cladograms were adjusted to fit vertical lines for easier interpretability and have no relative meaning; the phylogeny is purely to depict branching pattern and hypotheses tested. Red vertical lines in A and C and red data points in B and D indicate species hypothesis decisively favored by Bayes Factor analyses (see Supplemental Table 5 for values).

**Figure 4.**
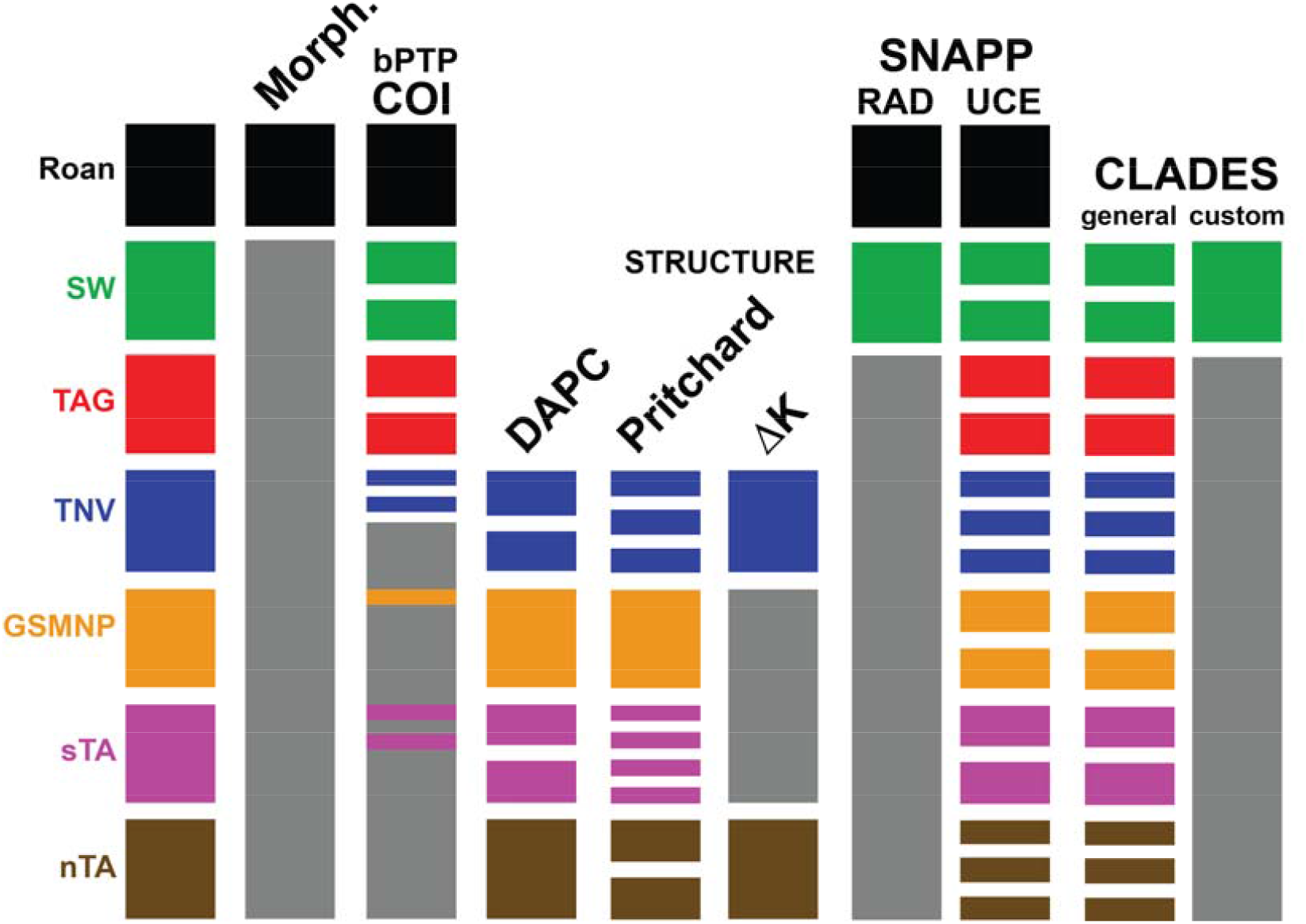
Species delimitation results supported across data and analysis types.

## Discussion

Major portions of the tree of life include branches (species) that are unknown or poorly known, and integrative studies in such taxa are challenging for many reasons. At the same time, collecting sub-genomic data for these taxa has become increasingly easy, leading to species delimitations founded largely, if not entirely, upon genetic data. Many studies have documented overestimation of species numbers by MSC-based delimitation methods both in empirical (e.g., Neimiller et al. 2012; Satler et al. 2013; Barley et al. 2013; Hedin et al. 2015; Hedin and McCormack 2017; Huang 2018; Yang et al. 2019; Derkarabetian et al. 2019; Chambers and Hillis 2019), and theoretical / simulation studies (Barley et al. 2017; Sukumaran and Knowles 2017; Mason et al. 2020; Sukumaran et al. 2021). Increasing the number of loci may also increase support for an incorrect hypothesis in Bayesian analyses (Yang and Zhu, 2018), and most relevant here, supporting species level divergences for intraspecific populations (Huang 2018). While MSC-based analyses are perhaps the most popular, other methods have also been shown to overestimate species numbers in low-dispersal taxa (e.g., Fernández and Giribet 2014; Hedin et al. 2015). Researchers studying cryptic species with inherently high population structure and allo- or parapatric distributions face a dilemma when delimiting species. Current phylogenomic species delimitation analyses overestimate species numbers, and without morphology, behavior, or distribution to assist, the degree of over-splitting is unknown. Systematists researching these taxa must be conservative in final species hypotheses (e.g., Barley et al. 2018), while simultaneously acknowledging that the actual number of species is still underestimated.

In our study, most genetic analyses recover at least six lineages as species (Fig. 4), but the consensus favors three species (Roan, SW, all others), the latter two of which are morphologically cryptic. Most importantly, the supervised machine learning approach we employed with a “custom” training data set provided a reasonable and biologically informed hypothesis of three total species. Phylogenies derived from ddRAD and UCEs were essentially congruent, but the inferred number of species in the BFD* analyses differed. Both had a peak at K = 3, but UCE data had a higher likelihood at K = 15, and leading to this maximum we argue the MSC over-splits because these taxa clearly violate the assumption of panmixia within species. BFD* analyses using ddRAD data do not overestimate species. However, like UCE analyses, there is an obvious increasing trend in likelihood with increasing species numbers leading us to ask if K=3 would still be the most likely hypothesis if more sequenced samples from different collecting localities (i.e., populations) had been included. As a validation approach, only a limited set of species-level hypotheses are tested in empirical studies using BFD*. It might be beneficial for researchers using this approach, especially for low dispersal taxa, to fully explore hypotheses to see if “false positives” are more prevalent.

### Training Data Justification and Considerations

In this study we present a first attempt at demonstrating the potential of a supervised machine learning approach to genetic species delimitation using biologically relevant customized training data sets. Here we provide some justification for our training data set choice and considerations for future work. In the case of Opiliones, which are largely understudied from a modern genetic perspective, there is scant genome-scale species level data available to serve as training data. Many Opiliones studies focusing on species delimitation using genetic data identify or conclude the presence of cryptic species, resulting in uncertainty across species boundaries (e.g., Boyer et al. 2007; Fernández and Giribet 2014; Hedin and McCormack 2017). The level of congruence and support for species delimitations seen in *Metanonychus* is exceptional (Derkarabetian et al. 2019), making this the most suitable training dataset for other low-dispersal Opiliones. As such, this training dataset has the most potential for application to the many unresolved putative cryptic species complexes in Opiliones.

More specifically, genetic statistics can be useful in identifying the overlap of actual and potential species boundaries between data sets. For UCE loci, the mean K2P-corrected genetic distance across all *T. brunneus* samples is 2.98% (range 0.37–8.92) and falls within the range of genetic distances seen across species of *Metanonychus* (mean = 14.746, range = 1.59–26.91). Speciation events within *Metanonychus* span recent and older divergences, where the mean K2P-corrected divergence of COI across the shallowest species-level split is 13.15% and the deepest split has a mean divergence of 27.15%. Mean COI divergence across *Theromaster* samples is 6.51%, with a maximum of 20.2%. Taken together, these genetic measures indicate that any potential species-level divergences in *Theromaster* fall within the range of actual species divergences seen in *Metanonychus*. Sensitivity to the underlying model is an obvious and related consideration. In this regard, in taxa with better representation of genome-scale species level data, it would be worthwhile to explore varying combinations of suitable taxa in the training data set. For example, including shallower or deeper species divergences, or data from a phylogenetically more diverse range of taxa (i.e., from other genera or families with similar biological characteristics).

One concern relates to the differences in dispersal dynamics between the regions each taxon is found in (Pacific Northwest for *Metanonychus* and southern Appalachians for *Theromaster*). Dispersal dynamics and bio-/phylogeographic histories are different across these regions, where geologic and other abiotic factors (e.g., river formation) can drive the speciation process, especially for the ancient and more topographically complex southern Appalachian Mountains. Despite these differences, given the biological and ecological similarity of these taxa, we hypothesize that, while the dynamics differ across regions, their *responses* and associated genetic signatures to any abiotic factors effecting speciation will be similar. It follows that using taxa that inhabit the same region should increase the effectiveness of our supervised approach, much in the same way as comparative phylogeography is a powerful approach for elucidating common underlying regional biogeographic patterns (e.g., Carnaval et al. 2014; Espíndola et al. 2016).

There are many questions in relation to the choice of training and testing datasets that deserve further attention. How closely related must the taxa be to be considered similar enough for this approach? What is an appropriate divergence date threshold? How ecologically similar (e.g., degree of climatic variable overlap) should they be? Questions relating to similarity and divergence dates can be explored with further data sets, as well as the relative importance of genetic versus ecological similarity between training and testing taxa. This choice will in fact be dependent on the organismal type and may ultimately need to be somewhat subjective, where the experience and organismal knowledge of the taxonomist will be critical in determining suitability of any training data set. Niche overlap can be quantified, however, in the case of our study the differences in local climates and the geographic distance between their respective distributions makes assessing niche overlap difficult. Further, the bioclimatic variables used in species distribution modelling do not necessarily capture the similarity in microhabitat “climate” for taxa found underneath the forest surface, living in deep leaf litter and underneath woody debris. Future studies should advance towards quantifiable metrics that determine if a species group(s) is an appropriate training dataset, as well as attempting this approach on taxa that are more directly linked to the bioclimatic variables used in species distribution modelling (e.g., plants).

### Putting Biology Back into Genetic Species Delimitation

Genetic species delimitation is driven largely by computational tools, model testing, and a desire for objectivity in analyses. However, this is increasingly at the expense of considering the biological characteristics of organisms, and importantly, with little discussion of how to adequately resolve the negative effects resulting from improper fit of variable biological characteristics with the models used. Cryptic species, and those taxa that undergo non-ecological speciation, are one of the biggest challenges in species delimitation, as many species criteria cannot be used in practice. Moving forward, addressing the cryptic species conundrum with genetic data might best be approached by using information already available, in this case inferring species in difficult unknown systems by using data from robustly known taxa with similar biological characteristics and modes of speciation. Recent integration of machine learning in species delimitation (Pei et al. 2018; Derkarabetian et al. 2019; Smith and Carstens 2020) provides algorithmic options which are versatile and customizable. Here we used a system-specific “custom” training dataset in a supervised machine learning framework to delimit cryptic species, where the training data were derived from *Metanonychus*, a previously studied system with robust species supported through multiple data types and with similar biological characteristics to *Theromaster*. These taxa share the same biological and ecological characteristics, most importantly dispersal ability and microhabitat preference, and both undergo the same type of speciation, and as such are expected to have very similar underlying genetic patterns associated with populations and species.

The power of applying a supervised machine learning approach derives from the ability to create custom training data sets that are specific to each study system, and to various classes of genetic data (e.g., UCE, RADseq, Sanger), capturing the inherently different characteristics of genetic data types. In this way, our approach combines a computational tool ideally suited for species delimitation, in this case, a supervised machine learning algorithm as a classification tool, with our knowledge of the biology and natural history of the focal organisms derived from our organismal expertise, leading to more informed, relevant, and reasonable species delimitation decisions when relying on genetic data only. The recently developed program DELINEATE (Sukumaran et al. 2021) takes a similar approach, using the information from known species to calculate speciation parameters which are then applied to delimit unknown samples. Similarly, the reference-based taxonomic approach used to delimit putative new species based on genetic distances of known species in a group of closely related and ecologically similar lizards is promising (Leaché et al. 2021).

There are extremely well-studied systems for which genetic species delimitation is largely successful, for example the model organism *Drosophila* (Campillo et al. 2020). However, for poorly studied groups (perhaps the majority of life) where basic biological and ecological knowledge can be difficult or impossible to acquire, inferring any biological details in an unknown or new taxon commonly relies on generalization from similar or closely related taxa where that information happens to be known. Our approach using known species limits in a supervised machine learning framework to infer unknown limits is a logical analytical extension of this inference process and is universally applicable to species delimitation in any taxon, especially for cryptic species.

## Materials and Methods

### Study System and Taxon Sampling

The genus *Theromaster* Briggs, 1969 currently includes two described species: the widespread *T. brunneus* (Banks 1902), and the poorly described *T. archeri* (Goodnight and Goodnight 1942) from several caves in extreme northeastern Alabama. Because *Theromaster* are small (body length usually <3 mm), short-legged, and most often found in sheltered microhabitats under rocks and logs, we anticipate high levels of population genetic structuring and potential cryptic diversification in this genus, as seen in many other northern temperate Opiliones with similar biology (e.g., Hedin and Thomas 2010; Hedin and McCormack 2017; Derkarabetian et al. 2016; Derkarabetian et al. 2019). Cryptic diversification is further expected because *T. brunneus* occurs in one of the most topographically diverse and biodiversity-rich regions of North America, the southern Appalachian Mountains. The Southern Appalachians are a well-known biodiversity hotspot for animals including vertebrates (e.g., Crespi et al. 2003; Weisrock and Larson 2006; Kozak and Wiens 2010) and arthropods (e.g., Hedin 1997; Marek 2010; Hedin and Thomas 2010; Keith and Hedin 2012; Garrick et al. 2018; Caterino and Langton-Myers 2019). For arachnids, almost all available molecular datasets for “wide-ranging” taxa in this region indicate *in situ* phylogeographic diversification, with multiple lineages that likely represent cryptic species (e.g., summarized in Keith and Hedin 2012; Hedin and McCormack 2017; Newton et al. 2020).

### Theromaster brunneus

is known from less than 10 literature records (Goodnight and Goodnight 1942; 1960, Briggs 1969; Kury 2003) from western North Carolina, eastern Tennessee, northern Georgia, and northern Alabama. However, our own collections and museum specimens indicate a broader distribution (Figure 1) that is atypically large for a single species of northern temperate laniatorean Opiliones. We first constructed a distribution map for *T. brunneus*, based on original collections from the Hedin lab and collections of Dr. William Shear (specimens now housed at San Diego State University). Locality data for all specimens housed in the San Diego State Terrestrial Arthropod Collection (with SDSU_TAC or SDSU_OP catalog numbers) have been deposited at the Symbiota Collections of Arthropods Network (http://symbiota4.acis.ufl.edu/scan/portal/index.php). Our specimen sample spans the known geographic distribution of the genus, including specimens from near the type locality of *T. brunneus* (“valley of Black Mountains”, North Carolina). All samples used in this study are identified as or considered *T. brunneus*; specimens morphologically identifiable as the questionable *T. archeri* could not be collected. UCE-based phylogenomic analyses of travunioid harvestmen strongly support a *Theromaster* + *Erebomaster* clade (Derkarabetian et al. 2018); as such we used *Erebomaster* samples as outgroups for mitochondrial and UCE datasets, and rooted ddRAD phylogenies (without *Erebomaster*) based on the UCE topology.

### Mitochondrial Data Collection and Analysis

A partial fragment of the mitochondrial cytochrome c oxidase subunit I (COI) gene was amplified using PCR primers and conditions as in Hedin and Thomas (2010) and Derkarabetian et al. (2010) for a total of 39 *T. brunneus* samples from throughout its distribution. PCR products were purified using Millipore plates, Sanger sequenced in both directions at Macrogen USA (Rockville, MD), then edited and aligned manually using Geneious 10.1 (Biomatters Ltd.). Gene trees were reconstructed using RAxML v8 (Stamatakis 2014), with a GTR GAMMA model applied to separate codon partitions. RAxML was called as follows: -# 500, -n MultipleOriginal, -# 1000, -n MultipleBootstrap. The resulting COI RAxML gene tree was used as input for bPTP species delimitation analyses through the bPTP server (https://species.h-its.org.ptp; Zhang et al. 2013). Two replicate analyses were run for 100,000 generations, with thinning at 100, and a burnin of 0.1.

### Morphology

Adult male *T. brunneus* have distinctive cheliceral modifications, making this taxon easily recognizable. Whole specimen and cheliceral segment digital images were captured using a Visionary Digital BK plus system (http://www.visionarydigital.com). Multiple individual images were merged into a composite image using Helicon Focus 6.2.2 software (http://www.heliconsoft.com/heliconfocus.html). For imaging, left chelicerae and pedipalps were dissected from male specimens. Male penises that were not already protruding from the genital operculum were physically extracted using a blunt insect micro pin. Chelicerae, penis, and pedipalps were examined using scanning electron microscopy (SEM). Specimens destined for SEM imaging were mounted onto stubs, critical point dried, coated with 6 nm platinum, and imaged on the FEI Quanta 450 FEG environmental SEM at the San Diego State University Electron Microscope Facility.

### ddRADSeq Data Collection and Analysis

Sixty-one *Theromaster* specimens from 50 distinct geographic locations were included in a “complete” (n=61 samples) matrix for ddRAD analyses (Fig. 1). Preparation of the ddRAD libraries followed Burns et al. (2016) and Derkarabetian et al. (2016), adapted from Peterson et al. (2012). DNA was extracted from whole specimens using the Qiagen DNeasy Blood and Tissue Kit (Qiagen, Valencia, CA) following the manufacturer’s protocol. We used restriction digest enzymes *EcoRI*-HF and *Msp1* (New England Biolabs, Ipswich, MA), and the corresponding adapters from Peterson et al. (2012). Briefly, ∼500 ng of genomic DNA was digested for 3 hours in a 50 μl reaction with 100 units each of the restriction enzymes *EcoRI*-HF and *Msp1* (New England Biolabs, Ipswich, MA), and 1X CutSmart Buffer (New England Biolabs). Samples were purified using Agencourt AMPure XP bead cleanup (Beckman Coulter, Inc., Brea, CA). Adapters were ligated to digested DNA in a 40 μl reaction that consisted of 33 μl digested DNA, 1.05 μM *MspI* P2 adapter, 0.54 μM *EcoRI* P1, 400 units of T4 DNA-ligase, and 1X T4 DNA ligase reaction buffer (New England Biolabs). Ligation reactions were incubated at room temperature for 40 minutes, heat killed at 65°C for 10 minutes, then cooled to room temperature at a rate of 2°C per 90 seconds. Samples with different adapters were pooled by column and then purified using AMPure XP bead cleanup. Pooled samples were then size selected to a size range of 400–600 bp with a Pippen Prep automated size-selection instrument (Sage Science, Beverly, MA). Primers with Illumina indices were added to the pooled samples using PCR; 50 μl reactions consisted of 23 μl DNA template, 2 uM PCR Primer P1, 2 uM PCR primer P2 (eight types for second-tier multiplexing, one per pooled sample), and 1X Phusion High Fidelity PCR Mastermix (New England Biolabs). Cycle conditions were 98°C for 30 seconds, 12 iterations of 98°C for 10 seconds and 72°C for 20 seconds (with a 16% ramp to slow cooling), and 72°C for 10 minutes. PCR products were purified via AMPure XP bead cleanup and quantified using a Bioanalyzer (Agilent Technologies, Santa Clara, CA). A pool consisting of an equimolar quantity of each library was sequenced as 100 bp SE reads on the Illumina HiSeq2500 platform at the University of California, Riverside’s Institute for Integrative Genomics Biology – Genomics Core Facility.

ddRAD data were processed using the *denovo* assembly method of ipyrad v.0.5.15 (Eaton and Overcast 2020), with the following settings adjusted from default: mindepth_majrule = 6, clust_threshold = 0.9, filter_adapters = 2, filter_min_trim_len = 35, max_Indels_locus = 4, max_shared_Hs_locus = 0.1. For the full 61-sample dataset, we ran min_samples_locus at 31 (50% complete matrix, called 61_31) and 48 (∼75% complete matrix, called 61_48). Maximum likelihood analyses of 61_31 and 61_48 matrices were run with RAxML v8 (Stamatakis 2014) using 1000 rapid bootstrap replicates and the GTRGAMMA model. These analyses of 61-sample matrices indicated three divergent and early-diverging lineages (see Results). Given the congruent recovery of three early-diverging lineages, we excluded these early-diverging samples and re-ran min_samples_locus at 45 and 51 for a reduced 51-sample dataset (called 51_45 and 51_51 respectively). These datasets only included “Tennessee Valley” (TNV), Great Smokey Mountain National Park (GSMNP), “northern Trans Asheville” (nTA), and “southern Trans Asheville” (sTA) lineages. This strategy effectively increases the number of loci retained for these derived lineages (see Bryson et al. 2016).

Using unlinked SNPs (a single randomly sampled SNP per locus) from the 61_48 matrix, we reconstructed both a “lineage tree” (individuals as OTUs) and “species tree” using SVDquartets (Chifman and Kubatko 2014) in PAUP* v4.0a152 (Swofford 2003) with n = 500 bootstraps. For the species tree, specimens were partitioned into groups following major clades recovered in RAxML and SVDquartets lineage tree analyses. Using the 51_45 unlinked SNPs matrix (n=1122 unlinked SNPs), we performed k-means clustering of PCA-transformed data using the find.clusters R function (“adegenet” package) (Jombart 2008; Jombaet and Ahmed 2011). Missing data were replaced by the mean frequency of the haplotype in the sample (scaleGen(data, NA.method="mean”). The Bayesian information criterion (BIC) was used to compare clustering models with a maximum of K = 20, retaining all principal components, and replicating the analysis 10 times. We then conducted a discriminant analysis of principal components (DAPC), retaining approximately one-quarter of the principal components and all discriminant functions. Using the same unlinked SNPs matrix, STRUCTURE 2.3.4 (Pritchard et al. 2000) runs were conducted using an admixture model with uncorrelated allele frequencies. All other priors were left as default. For individual K values ranging from 2–12, analyses were replicated four times, each run including 200,000 generations with the first 20,000 generations removed as burnin. Data were summarized using CLUMPAK (Kopelman et al. 2015), with a best-fit K chosen utilizing the ΔK method of Evanno et al. (2005), and the prob (K) method of Pritchard et al. (2000).

Based on phylogenetic analysis of the UCE and 61-sample RADSeq datasets, plus STRUCTURE (Pritchard et al. 2000) and DAPC (Jombart et al. 2010) analyses of the 51-sample RADSeq dataset (see Results), we chose 18 samples to represent all primary *Theromaster* lineages. For these 18 samples we re-ran ipyrad using settings as above, at 18_12 occupancy. From this, an unlinked SNPs file was created using the Phrynomics package (Leaché et al. 2015; github.com/bbanbury/phrynomics), where nonbinary characters were removed and bases were translated. To further visualize genetic structure and clustering within *Theromaster*, we analyzed this dataset using a Variational Autoencoder (VAE) implemented with a modified version of the sp_deli script (https://github.com/sokrypton/sp_deli) derived from Derkarabetian et al (2019). The matrix was run through the VAE five times and the analysis with the lowest average loss after removing 50% burnin was considered the optimal output.

### UCE Data Collection and Analysis

Studies have shown the utility of UCEs at shallow levels (e.g., Smith et al. 2014; Zarza et al. 2016), particularly in arthropod taxa (e.g., Blaimer et al. 2016; Hedin et al. 2018; Derkarabetian et al. 2019). The UCEs targeted in the arachnid probe set (Faircloth 2017) are exonic in origin with the “core” UCE corresponding to coding region while the “flanking” region are non-coding introns (Hedin et al. 2019). As such, in taxa with high population structure, flanking non-coding regions of UCEs are informative for population level structure and can be used in phylogenomic species delimitation analyses (Derkarabetian et al. 2019). Representative samples for UCE sequencing were chosen based on preliminary RAD analyses, subsampling main lineages (See Figure 2).

Sequence capture of UCEs followed the protocol of Derkarabetian et al. (2018). A subset of 15 samples representing all major ddRAD lineages were used in UCE experiments and were prepared in multiple library preparation and sequencing experiments. Protocols across these experiments were largely identical, differing mainly in sequencing platform. Genomic DNA was extracted from either multiple legs or whole bodies using the Qiagen DNeasy Blood and Tissue Kit (Qiagen, Valencia, CA). Extractions were quantified using a Qubit Fluorometer (Life Technologies, Inc.) and quality was assessed via gel electrophoresis on a 1% agarose gel. Up to 500 ng were used in DNA fragmentation procedures, either using a Bioruptor or a Covaris M220 Focused-ultrasonicator, as in Derkarabetian et al. (2018). UCE libraries were prepared using the KAPA Hyper Prep Kit (Kapa Biosystems), using up to 250 ng DNA (i.e., half reaction of manufacturer’s protocol) as starting material. Ampure XP beads were used for all cleanup steps. For samples containing <250 ng total, all DNA was used in library preparation. Target enrichment was performed on pooled libraries using the MYbaits Arachnida 1.1K version 1 kit (Arbor Biosciences, Ann Arbor, MI) following the Target Enrichment of Illumina Libraries v. 1.5 protocol (http://ultraconserved.org/#protocols). Hybridization was conducted at either 60 or 65 °C for 24 hours, with a post-hybridization amplification of 18 cycles. Following an additional cleanup, libraries were quantified using a Qubit fluorometer and equimolar mixes were prepared for sequencing either with an Illumina NextSeq (University of California, Riverside Institute for Integrative Genome Biology) with 150 bp PE reads, or an Illumina HiSeq 2500 (Brigham Young University DNA Sequencing Center) with 125 bp PE reads (see Suppl. Material 1).

Raw demultiplexed reads were processed with the Phyluce pipeline (Faircloth 2016). Quality control and adapter removal were conducted with the Illumiprocessor wrapper (Faircloth 2013). Assemblies were created with Velvet (Zerbino and Birney 2008) at default settings. Contigs were matched to probes using minimum coverage and minimum identity values of 65. UCE loci were aligned with MAFFT (Katoh and Standley 2013) and trimmed with Gblocks (Castresana 2000; Talavera and Castresana 2007) implemented in the Phyluce pipeline. All individual UCE loci were imported into Geneious 10.1 (Biomatters Ltd.) and manually inspected to check for obvious alignment errors and remove putatively non-homologous sequences (e.g., any sequences more divergent than outgroup taxa).

Concatenated and partitioned phylogenetic analyses were run on two datasets differing in the taxon coverage needed to include a locus in the final dataset: 50% and 70%. Maximum likelihood analyses were run with RAxML v8 (Stamatakis 2014) using 200 rapid bootstrap replicates and the GTRGAMMA model. Using the 70% concatenated UCE matrix we also reconstructed a lineage tree using SVDquartets (Chifman and Kubatko 2014) with n = 500 bootstraps. Finally, we made a 50% taxon coverage unlinked SNP dataset from alignments with a custom wrapper script using snp-sites (Page et al. 2016) to convert alignments to vcf format, randSNPs_from_vcf.pl (https://www.biostars.org/p/313701/) to select a single random SNP from each alignment’s vcf file, vcf2phylip.py (https://github.com/edgardomortiz/vcf2phylip) to convert vcf files back to phylip, and AMAS (Borowiec 2016) to concatenate all randomly selected SNPs into a single phylip file. The Phrynomics R package (Leaché et al. 2015; https://github.com/bbanbury/phrynomics) was used to select only biallelic SNPs and translate SNPs to integers. The VAE was run on this dataset as done with ddRAD data.

### Bayes Factor Delimitation* Analyses

We conducted BFD* (Grummer et al. 2014; Leaché et al. 2014) species delimitation analyses using SNPs derived from both the ddRAD and UCE data using SNAPP (Bryant et al. 2012) implemented in the BEAST 2.4.5 package (Bouckaert et al. 2014). Analyses were run on the 18_12 ddRAD and the 50% taxon coverage UCE datasets. For each SNP dataset we tested multiple alternative species hypotheses. Hypotheses tested were derived from other data types and analyses used in this study including morphological, COI, and phylogenetic and STRUCTURE/DAPC analyses of nuclear data. To test the BFD* approach to its fullest extent in this study system (and hence its potential to delimit populations), we also included a nested set of hypotheses up to the maximum potential number of species, where every specimen was considered a different species. All BFD* analyses were run for 100,000 generations, with 10,000 generations as pre-burnin, 48 steps, and an alpha value of 0.3. Two replicates of each analysis were run to check for convergence. A comparison of marginal likelihoods was conducted using Bayes factors (Kass and Raftery 1995), with values above 10 considered to be decisive support.

### Supervised Machine Learning Analyses

We analyzed UCE loci with the supervised machine learning species delimitation program CLADES (Pei et al. 2018). CLADES is a classification model derived from a type of machine learning algorithm called a support vector machine to classify samples as either “same species” or “different species” using multi-locus data in a two-species model. CLADES computes five summary statistics from the data (both training and testing) and uses these statistics as features to create the model and classify samples: private positions, folded-SFS with *k* bins, pairwise difference ratio, F_ST_, and longest shared tract (defined in Pei et al. 2018). A training data set, where pairwise comparisons of all samples are defined *a priori* as either the same or different, is used to build the model and classify a test dataset with unknown species status.

We analyzed *Theromaster* UCE data in CLADES using two training data sets. First, Pei et al. (2018) provided a training data set called “All” (which we refer to as “general” here) based on simulated data with varying values of theta (Θ), migration rate, and divergence time under a two species model. This “general” data set is meant to be broadly applicable across taxa as simulated data encompass the broad diversity of genetic patterns across plants and animals (Pei et al. 2018). Second, we developed a “custom” training data set derived from the well-known, robust species of *Metanonychus* recently revised in an integrative taxonomic context (Derkarabetian et al. 2019). All *Metanonychus* species are easily diagnosed based on both somatic and genitalic morphology, with both mitochondrial and nuclear data supporting species status across a broad array of analysis types. Most importantly, *Metanonychus* and *Theromaster* share similar biological and ecological characteristics, including low dispersal ability and microhabitat preferences. Microhabitat preference and ecological similarity are largely based on our experience collecting these taxa over many years. Quantifying biological and ecological similarity, as well as dispersal ability, is difficult in poorly-known taxa with cryptic biology, like those that occupy “hidden” microhabitats (see Discussion for further justification).

For both *Theromaster* and *Metanonychus*, datasets included all UCE loci shared across all samples in each data set (n= 52 for *Theromaster*, n = 12 for *Metanonychus*) (the “spp” dataset of *Metanonychus* in Derkarabetian et al. 2019). To create the “custom” *Metanonychus* training dataset, we ran the *Metanonychus* UCE loci through CLADES against the “general” training dataset. As expected, and found in Derkarabetian et al. (2019), analyses favored all populations as species, and all output files reflected this. The output files contain pairwise comparisons of all specified populations; those files with pairwise comparisons between populations that belonged to the same species as delimited in Derkarabetian et al. (2019) were manually modified to reflect that the samples belonged to the same species (switching +1 to -1). All relevant files required for the model (see CLADES documentation) were manually created from these output files. LibSVM (Chang and Lin 2011) was used to create the “*.model” file from the “*.sumstat.scale” file using default parameters, with the addition of training for probability estimates (-b 1). We excluded the Roan sample from CLADES analyses because this specimen (i.e., species) is morphologically distinguishable from the rest of *T. brunneus* and is represented by only a single specimen. In this way our goal was to assess the success of this approach on a dataset consisting only of morphologically similar lineages that are putative cryptic species.

## Acknowledgements

We would like to thank Derek Hennen, Matthew Niemiller, Bill Shear, and Kirk Zigler for providing important specimens. For assistance with fieldwork we thank Jason Bond, Fred Coyle, Ryan Faucett, Dalton Hedin, Lars Hedin, Robin Keith, and Steven Thomas. Erik Ciaccio and Morganne Sigismonti helped with specimen imaging. Ingrid R. Niesman provided SEM support at SDSU. This work was supported by the National Science Foundation (DEB 1354558 to M.H.).

## Notes

### Competing Interest Statement

The authors have declared no competing interest.

