## Supplemental Figures for "The challenge of delimiting cryptic species, and a supervised machine learning solution"

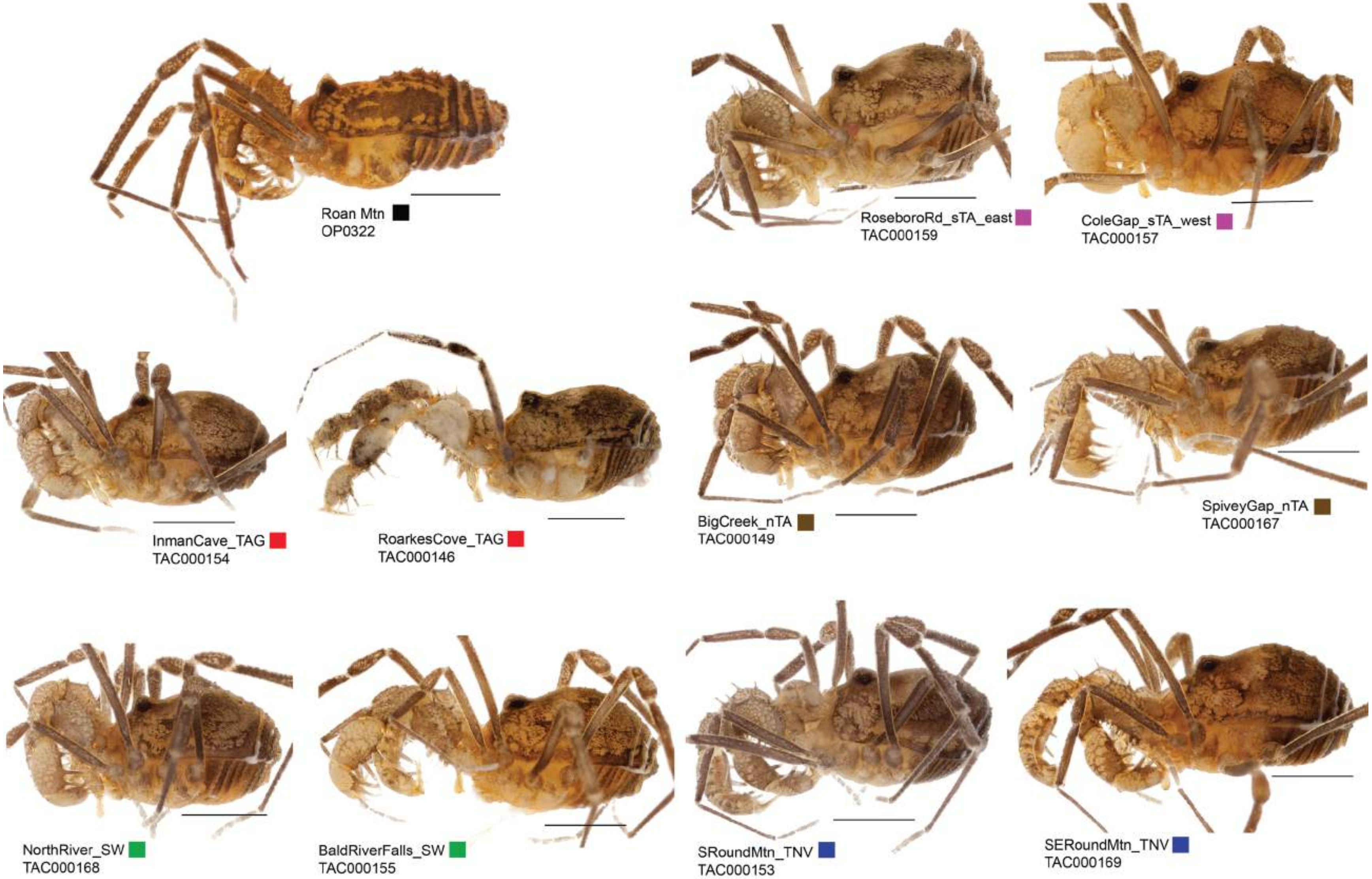


**Figure S1.** Habitus images of adult males. Colored squares correspond to primary phylogenomic lineages.

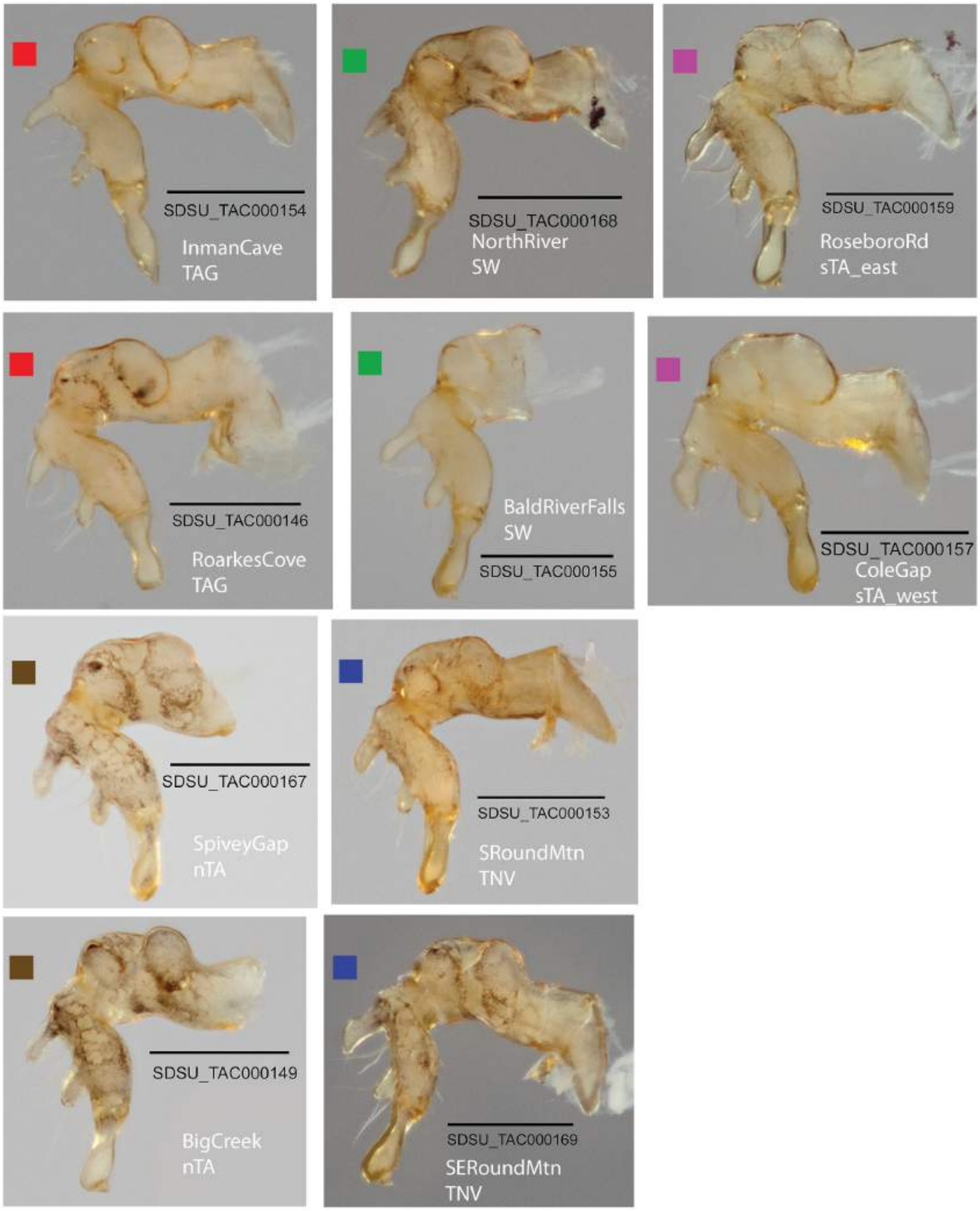


**Figure S2.** Digital images of adult male chelicerae. Colored squares correspond to primary phylogenomic lineages.

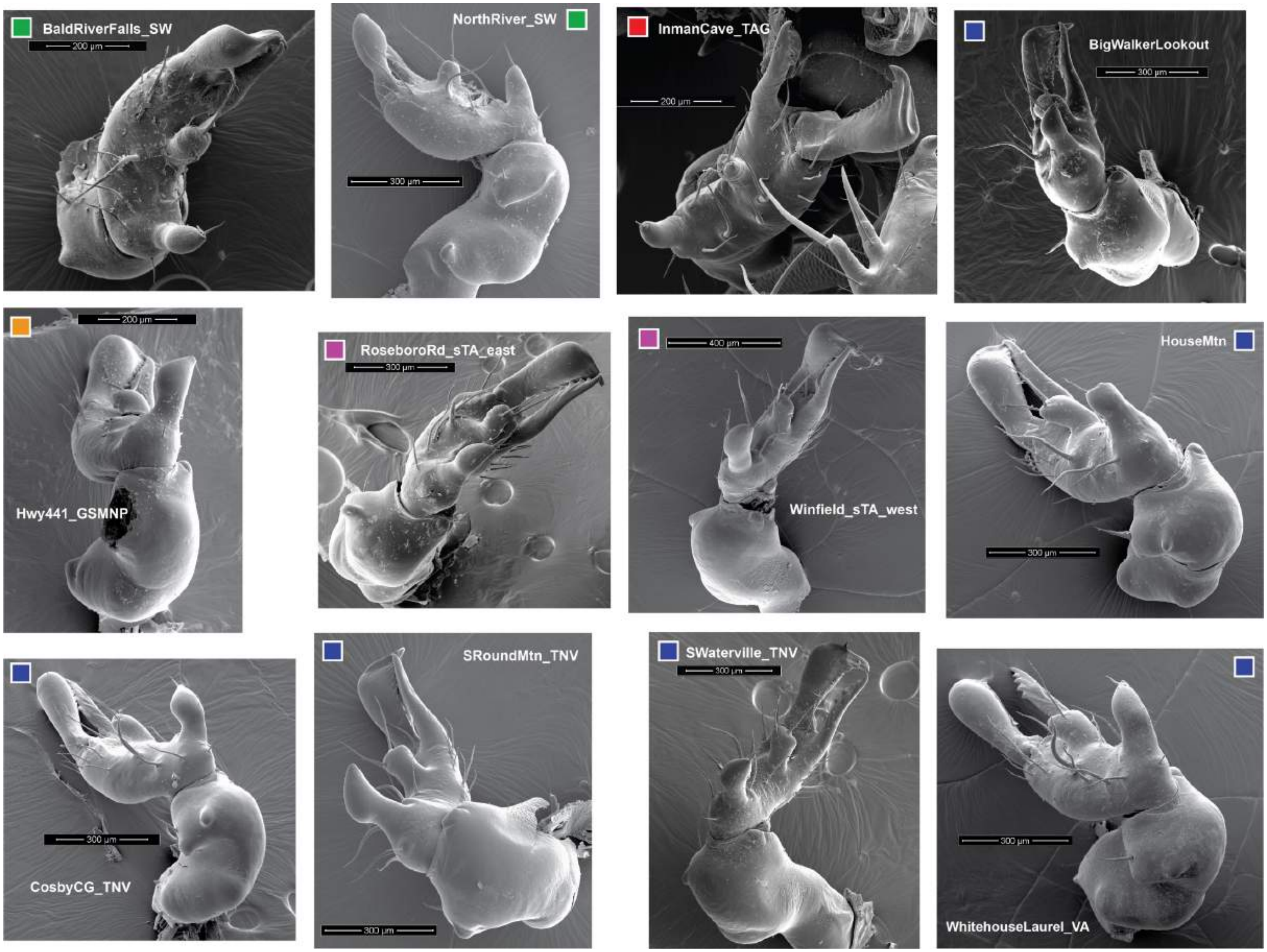


**Figure S3.** SEM images of adult male chelicerae. Colored squares correspond to primary phylogenomic lineages.

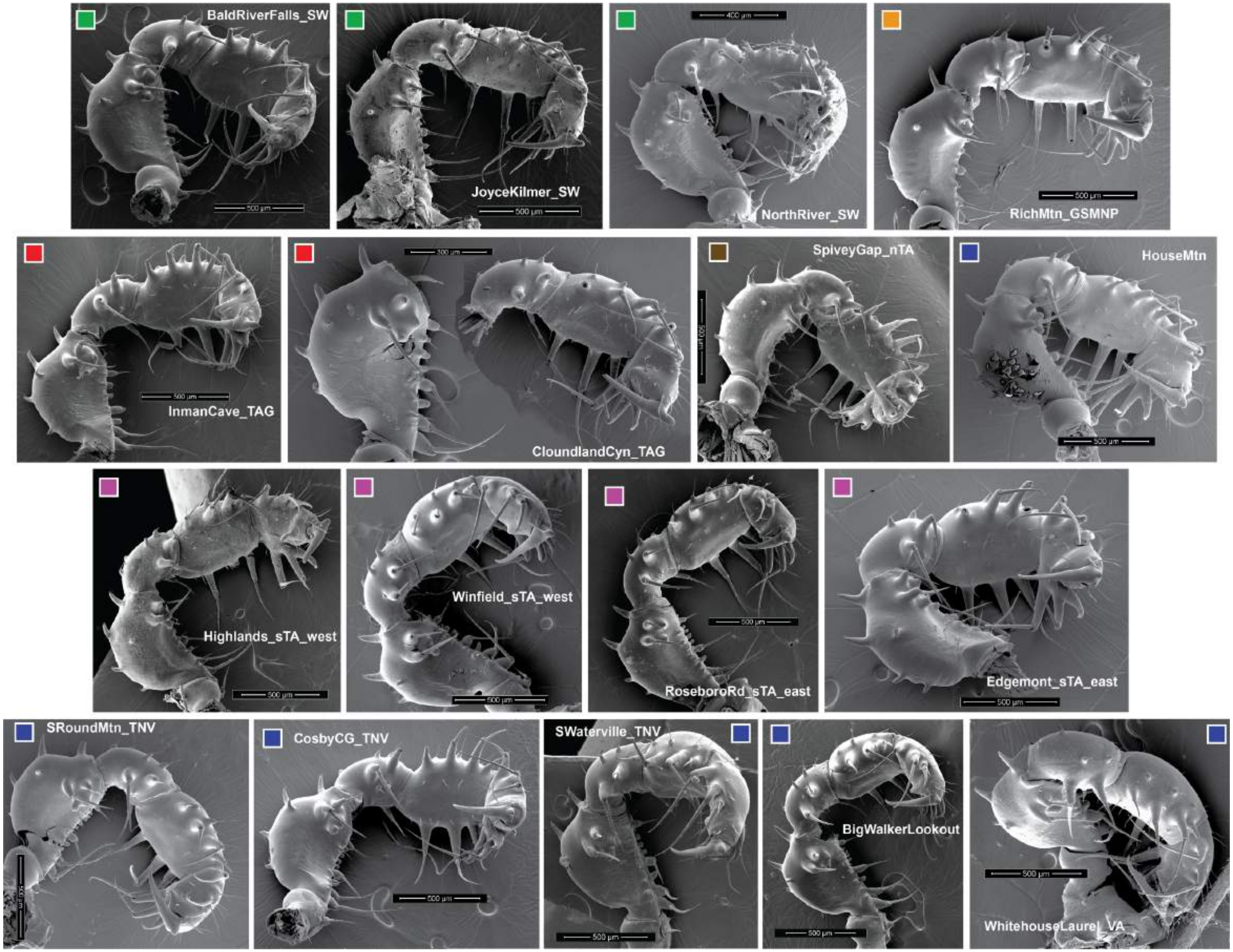


**Figure S4.** SEM images of adult male pedipalps. Colored squares correspond to primary phylogenomic lineages.

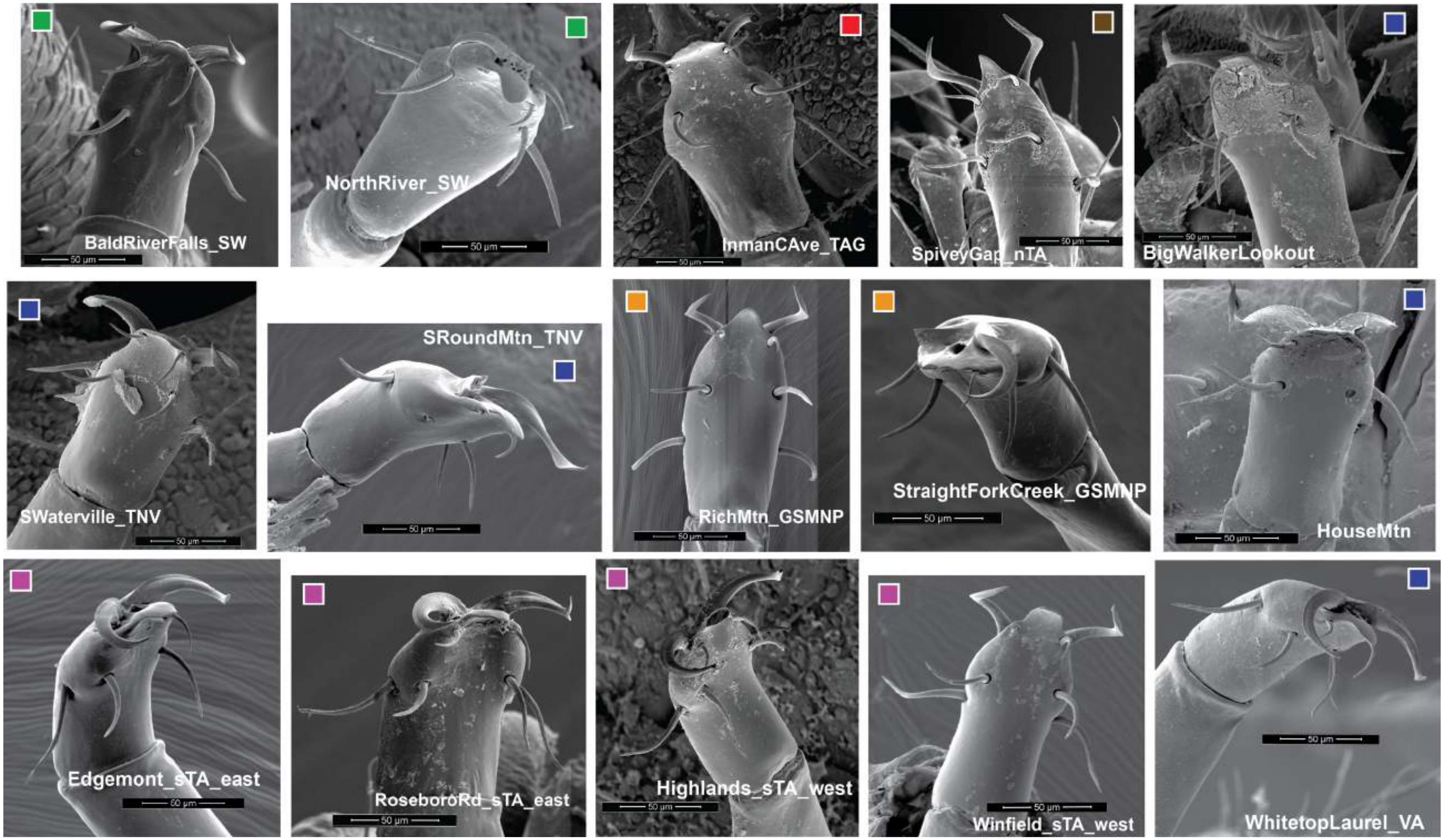


**Figure S5.** SEM images of adult male genitalia. Colored squares correspond to primary phylogenomic lineages.

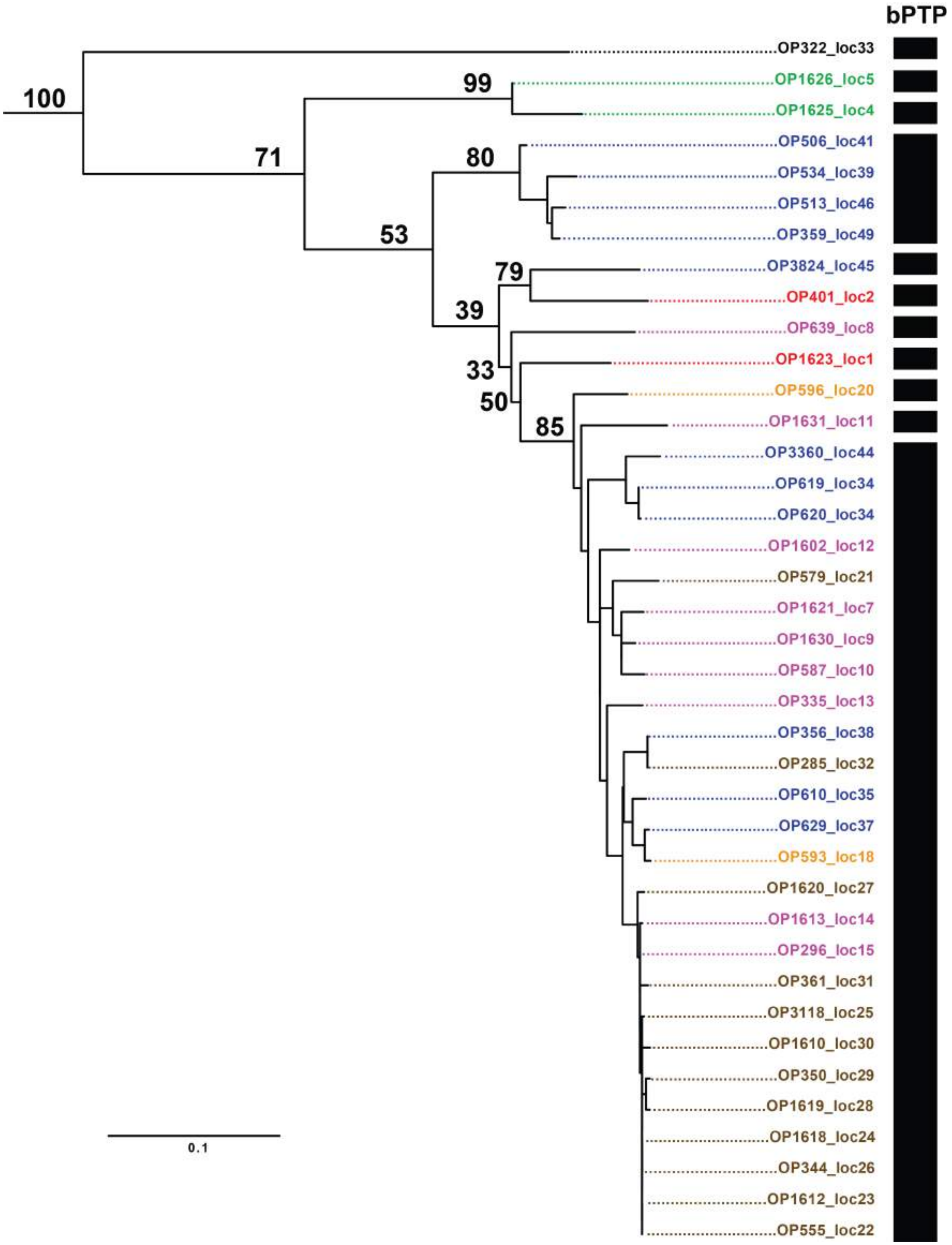


**Figure S6.** COI RAxML phylogeny. Bootstrap support indicated for major nodes only. Colors correspond to major phylogenomic lineages. Bars on the right indicate species supported by bPTP analyses.

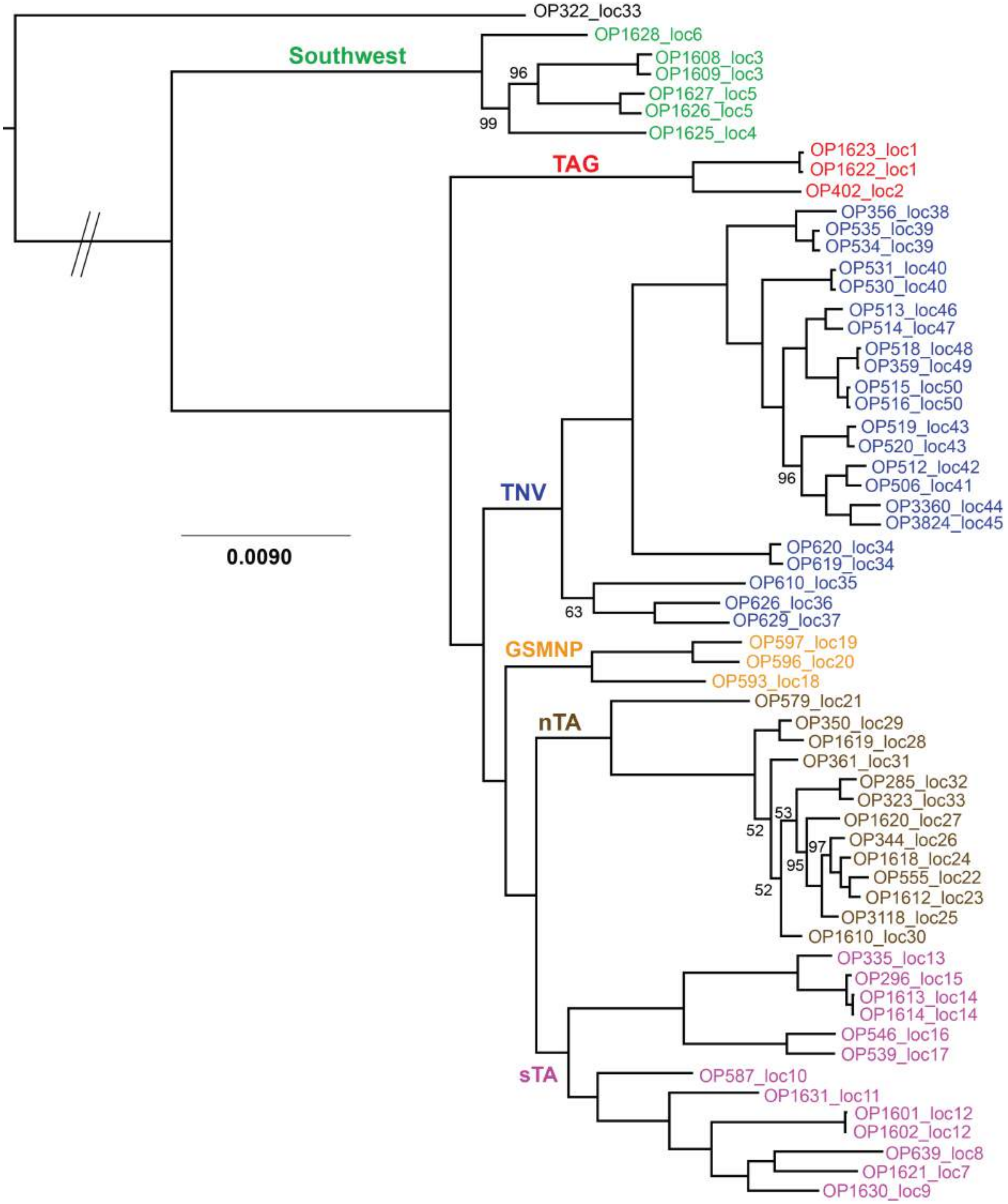


**Figure S7.** RAxML phylogeny of 61_31 ddRAD matrix (see Supplemental Material I for matrix details).

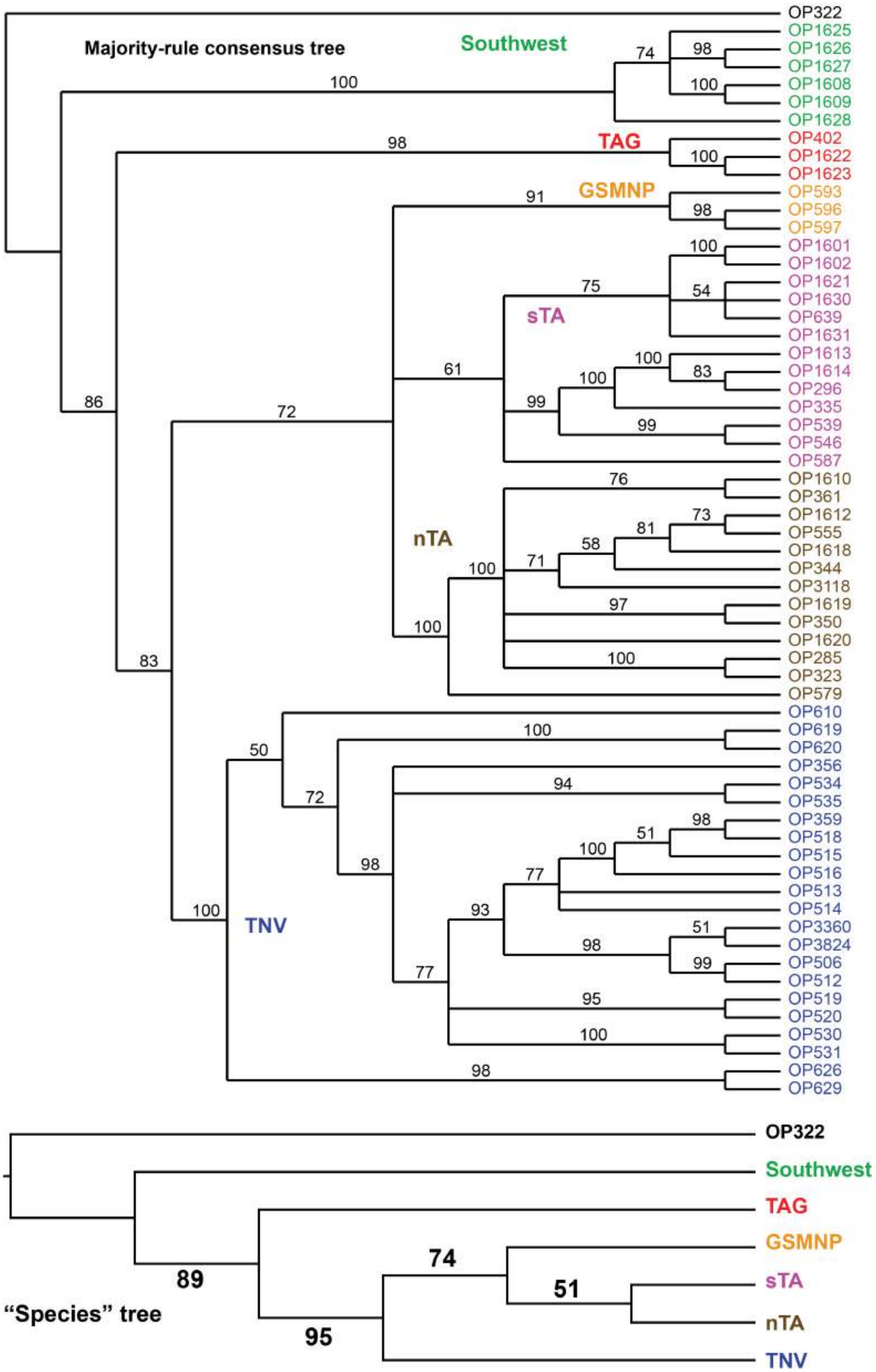


**Figure S8.** SVDQuartets phylogeny of ddRAD SNPs from 61_48 matrix for lineage tree (top) and species tree (bottom). See Supplemental Material I for ddRAD matrix details.

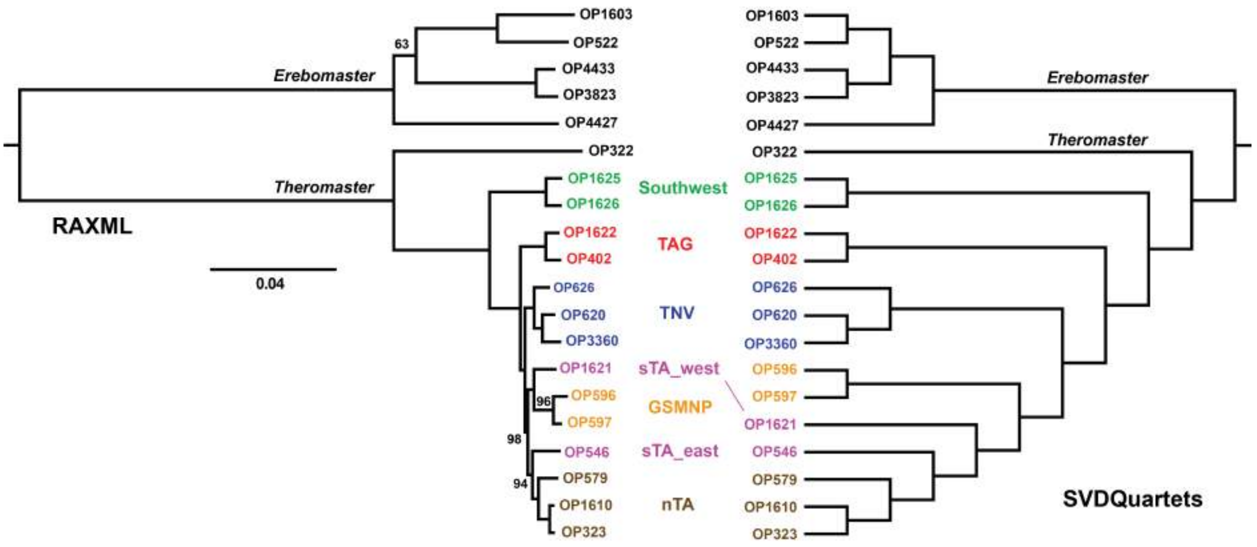


**Figure S9.** Phylogenomic analyses of UCE data using RAxML (left) and SVDQuartets (right) based on the 70% taxon occupancy matrix. For RAxML, all nodes have 100% bootstrap support unless indicated.

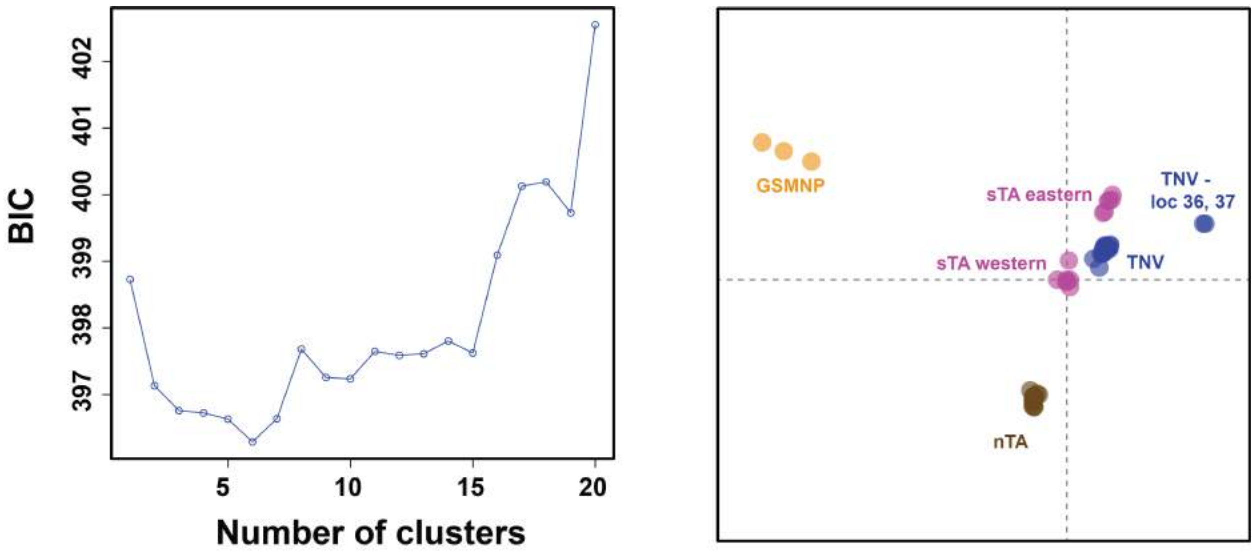


**Figure S10.** Results of DAPC analyses on 51_45 ddRAD SNP dataset (*see Supplemental Material 1 for matrix details).

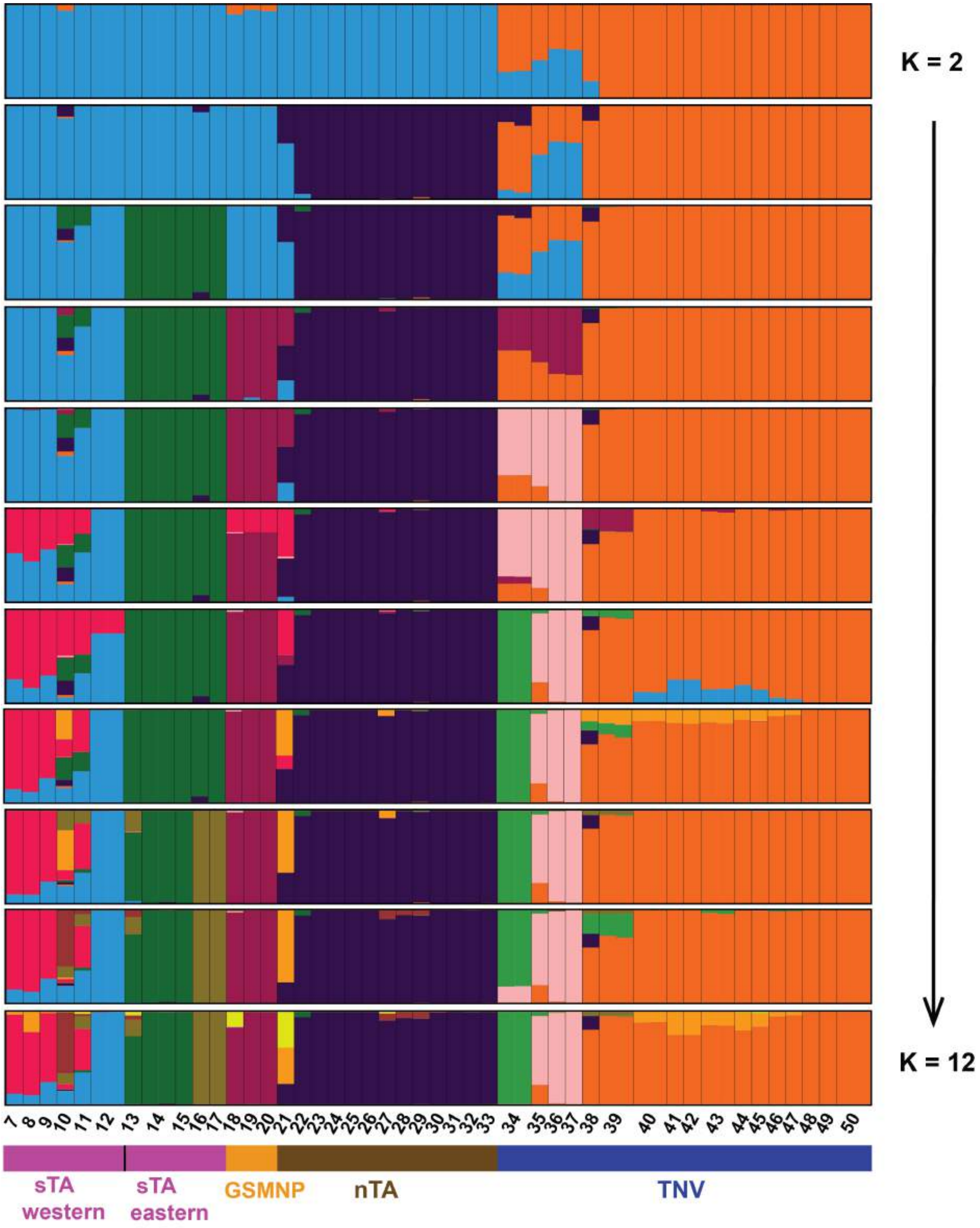


**Figure S11.** Results of STRUCTURE analyses on 51_45 ddRAD SNP dataset (*see Supplemental Material 1 for ddRAD matrix details).

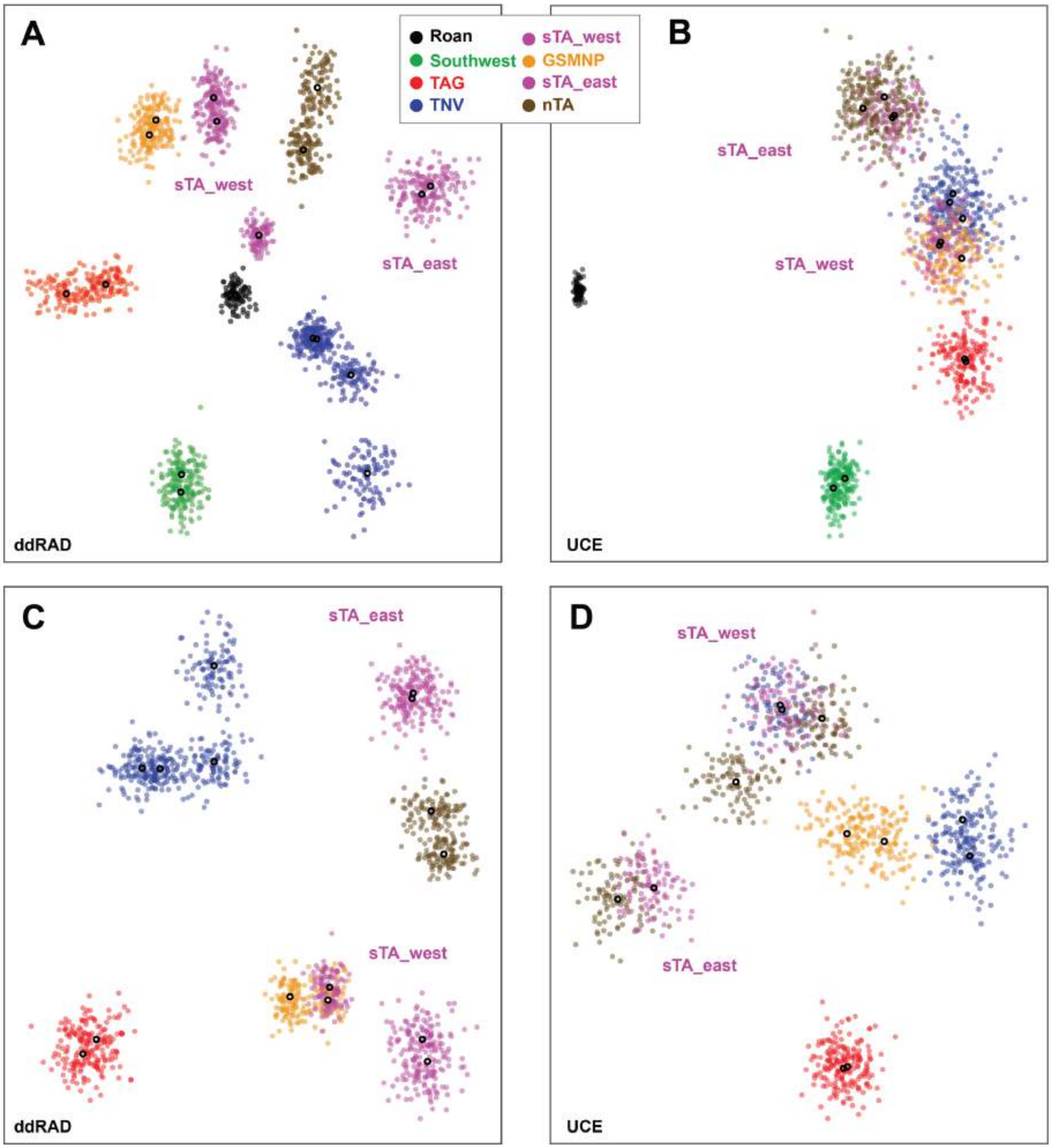


**Figure S12.** VAE plots for 51_45 ddRAD (A and C) and 50% taxon occupancy UCE (B and D) SNP datasets. See Supplemental Material 1 for matrix details. C and D exclude Roan Mountain and Southwest samples.
